# FUNCTIONAL AGEING: A Science-Technology-Society Approach to teach Ageing and Age-related Diseases

**DOI:** 10.1101/2023.05.25.538223

**Authors:** Gavin Ng, Vismitha Rajeev, Sharmelee Selvaraji, Karthik Mallilankaraman, Manoor Prakash Hande

## Abstract

**Purpose:** Ageing is a complex biological process that involves numerous genes and pathways. Experimental studies have identified several of these genes and pathways that that can extend or shorten an individual’s lifespan. Understanding these genes and pathways can help develop interventions that improve health and quality of life of older people. Nonetheless, as the global population continues to age, it is essential to comprehend the ageing process’s impact on society. Functional age is defined as a combination of chronological, biological, and psychological ages. An educational course called “Functional Ageing” was created for life science students at the National University of Singapore (NUS) during the academic year 2016-2017. The module adopted an interdisciplinary approach based on science-technology-society (STS) methodology and aimed to equip students with the analytical tools needed to assess the ageing process at both the molecular and physiological/functional levels. Ultimately, the module aimed to promote the understanding of ageing processes, particularly functional ageing in a population and its societal impacts.

**Methods:** This “Functional Ageing” course spanned over 13 weeks, consisting of weekly four-hour sessions that aimed to integrate both biology of ageing and societal perceptions of an ageing population. The first half of the semester covered the molecular processes that govern ageing, while the second half focussed on societal perception, burden of disease, healthy ageing interventions and creating an ageless society. Experimental and epidemiological studies were used to explain the ageing process. Expert guest lecturers were invited throughout the module to share their experiences on the demanding research areas of ageing today, as well as clinical aspects of age-related diseases. In addition, a field visit to a geriatric ward at the mental health organisation was arranged to showcase the country’s approach to dealing with the rising demands of ageing, and to provide experiential learning that can help inculcate a clearer public perception of ageing.

**Results:** This 13-week module helped to improve students’ perception about functional ageing in today’s society. After the module, a reflection analysis was conducted to evaluate students’ perceptions of ageing society. Overall findings garnered have demonstrated that while students generally have a brief understanding of the biological processes of ageing, their perception on how ageing is being manifested in a public health and societal setting is in paucity. However, the routine university-directed feedback survey indicated that the module had a positive impact on students’ appreciation and comprehension of the interplay between biological and sociological aspects of ageing, thus achieving the learning objectives of the module.

**Conclusion:** The aim of the “Functional ageing” module was to integrate the biology and sociology of ageing to provide a better interdisciplinary understanding of ageing in society. After completing the module, students demonstrated a change in their mindset and attitude towards how their scientific understanding of ageing can be coupled to their public perception of ageing. As ageing populations become more prevalent in many societies, it is crucial that we prepare for an ageing population and develop a positive perception of ageing. Through educational efforts like this course/module, students will exhibit greater awareness of ageing issues, leading to more informed and balanced perceptions.

## Background

“Nothing ever becomes real till it is experienced”. These words by John Keats are especially true in the realm of education. Science Technology Society (STS) learning approach is, as its name states, an integration of science, technology, and society. It allows students to learn the impact of science and technology on the society through real-life experiences^1,2^. The STS approach because of its interdisciplinary nature, not only enhances students’ learning, but it also ensures relatability and creates an impact that follows through for a lifetime of learning.

Apart from imparting methodological knowledge, educators must also encourage undergraduates to push beyond the boundaries of scientific accuracy. To become skilled science communicators, students need to extend their creativity and adaptability to transform complex science into an engaging and inspiring narrative. Making science more accessible to the public is crucial, and this process should commence in undergraduate education^3^. With increased emphasis on experiential learning and establishing connections with the real world, there is an increasing need to adopt the STS/interdisciplinary learning approach to enhance the learning experience of students in a wide range of aspects. The interconnectedness of the world through internet and social media has made knowledge acquisition much easier than before. However, it has also led to a surge of opinionated biases being circulated^4^. There is hence a need to challenge the stigma that arises from these opinionated biases. The major concern for the public health sector is the masking of factual information about diseases and health concerns in the name of myths. The recent COVID-19 pandemic and the myths surrounding it is an example of great relevance and recency. Education is a powerful tool that can mould impressionable minds and it is therefore important to strengthen the foundation of the future. Additionally, higher education should aim to equip all graduates with the skills required to distinguish facts from falsehoods. The interdisciplinary approach thus allows widening of perspectives, critical thinking, and the ability to delineate facts from false information.

As much as facts are crucial, being open-minded and garnering perspectives are pivotal in developing solutions to problems. The growing emphasis on interdisciplinary studies and thinking reflects a change in the educational focus^5^. The STS or interdisciplinary approach is essential, as it not only provides diverse perspectives on a specific issue but also facilitates collaboration and enables us to utilize readily available resources to resolve the problem at hand. By engaging with individuals from diverse backgrounds, learning from their problem-solving techniques, and brainstorming to apply unique approaches to the present issue, the STS and/or interdisciplinary approach proves to be indispensable.

Apart from incorporating interdisciplinary perspectives and fighting stigma surrounding public health topics, it is also crucial to simplify complex biological concepts and converse, to advocate the importance of concepts that are often undermined. One such concept is ‘Ageism’, which refers to the stereotypes and discrimination against people based on the traits of age. This term was initially coined in 1969 to describe the discrimination against elderly^6^. Living in an era of a rapid increase in the ageing population, ageing has become a more pertinent concept than ever before. It is predicted that by 2030, 1 in 6 individuals will be 60 years old and above and that by 2050 the number will double to 2.1 billion ^7^. With ageing, age-related diseases are certainly on the rise. This includes dementia, Parkinson’s disease, cancer, osteoporosis, diabetes mellitus, hypertension, arthritis, and many others. Due to improved healthcare systems and technology, we now live longer than we previously could. This increase in longevity, unfortunately, comes with the cost of compromised health span. Therefore, there are various means by which the problem of ageing is being tackled at different levels of society. This includes policies favouring active ageing, greater insurance coverage for age-related diseases, creating awareness amongst public to age gracefully and preventive measures in the form of advocating lifestyle-associated changes and even supplementary pills ^8^. In the context of ageing, even the smallest action can have a domino effect. While these concepts can be theoretically explained and supported by statistics and facts, their true impact can only be experienced on the ground through observing, listening to, and empathizing with those who are directly affected by the circumstances. Hence, there is a need to educate and allow the younger generation to understand and take a hands-on approach in tackling the problem of ageing – be it at home or in the larger society. The “Functional Ageing” module/course for year 4 life sciences undergraduates at the National University of Singapore has thus taken proactive steps in inculcating the importance of understanding the concept of ageing in a holistic manner with the incorporation of the STS approach.

### Objectives and Study Plan

The introduction of ageing and its societal implications were integrated into a semester-long module at the National University of Singapore. This **“**Functional Ageing” module comprised of weekly four-hour sessions for 13 weeks. In this module, students were exposed to concepts and ideologies pertaining to ageing and society, with the aim of fulfilling the interdisciplinary nature of the topic. The course challenged students to recognize the significance of ageing in both scientific and public spheres. During the first half of the semester, the course covered the molecular processes underlying ageing, while the second half focused on societal perceptions, disease burden, interventions for healthy ageing, and creating an ageless society. The ageing process was explained based on experimental and epidemiological studies. What sets this module apart from other courses on ageing or public health is the adoption of both STS and science communication pedagogies with an interdisciplinary approach. While we have recently documented another course on Radiation and Society^9^ that adopts a similar approach for undergraduate students from diverse backgrounds, the current study plan is unique because it is designed for life science students with an integrative sciences or STS approach.

This module is conducted one semester per academic year, with each class size averaging 50 or less students. Offered as part of an undergraduate curriculum, it enrols students from life sciences discipline. While this is an elective module, it has a prerequisite of basic cell biology and ageing. Thus far, six cohorts of students (2016-2023) with a total strength of 268 have enrolled into this module.

During the introduction of this module, the learning objectives were communicated clearly to the students **(Complete syllabus not shown but can be provided upon request)**. At the end of this module, students will be able to:

- Learn about molecular processes governing the ageing process;
- Learn more about genetics and epigenetics of ageing;
- Gain knowledge on developing therapeutic strategies of ageing and age-associated disorders;
- Learn about translational science of ageing;
- Compare and contrast the processes of ageing and cancer – relevance to other age-related diseases;
- Critically evaluate the social perception of ageing and its impact on society;
- Assess and evaluate the biological, social and psychological aspects of ageing;
- Understand the concepts of healthy ageing and an age-less society.

Throughout the course, experts either directly or indirectly associated with ageing issues delivered lectures on recent developments and their impacts on today’s society. Notably, Professor Kua Ee Hock, Department of Psychological Medicine, National University of Singapore discussed the cohort studies on dementia management using interventions such as music therapy and art therapy in Singaporean elderly population (Academic Year 2016 -2017, **Figure 1a**). In the most recent version of the module, Professor Leonard Hayflick, known for his discovery on the Hayflick Limit, was invited to share his research findings on ageing and discuss how the perception of ageing has evolved over the long term with the students. (**Figure 1b**).

**Figure 1.** Guest Lectures for the module. (a) Professor Kua Ee Hock, Department of Psychological Medicine, National University of Singapore discussed about interventions of art and music for dementia management in elderly population of Singapore (Academic Year 2016-2017). (b) Professor Leonard Hayflick (well known for his discovery of “Hayflick Limit”) Department of Anatomy, University of California San Francisco, USA was invited as a guest speaker to share with students enrolled into this module about his research experiences and his motivation in engaging ageing research (Academic Year 2021-2022)

Other notable topics covered in this module includes:

1. Theories of ageing
2. Telomeres and DNA damage theory of Ageing and others
3. Age related diseases
4. Epidemiology of Ageing – Lifestyle factors (Diabetes etc.)
5. Evolution and Ageing; Ageing and Cancer
6. Determinants of Health-Span
7. Interventions – calorie restriction and exercise
8. Ageing Society – or Age-less society
9. Life span, health-span and demography of ageing
10. Ageing and quality of life – cognitive decline, dementia, frailty, sarcopenia
11. End-of-life challenges and long-term healthcare
12. Societal challenges in the ageing population
13. Future of ageing

As part of the module, students were taken on a field visit to a local health institute to gain insight into how the healthcare system is being restructured to meet the increasing demands of an ageing population. Professionals working at the organization shared their perspectives with the students, providing them with a better understanding of ageing issues and debunking common misconceptions. **(Figure 2)**. During the visit, students had the opportunity to observe how mild-dementia patients were cared for in the geriatric wards of the centre. In earlier iterations of the course, students witnessed how intervention studies for dementia, such as music therapy and art therapy sessions, were implemented. Furthermore, to ensure that students were kept abreast of their learning, a mid-term assessment was conducted to examine on topics taught during lectures. As part of the assessment, students were required to write a reflective essay to stimulate their thinking on what they had learned during field trips and to prompt them to develop their perspectives on how ageing impacts society. Students were encouraged to draw on their experiences from visiting the eldercare institute, as well as readings and lectures. Additionally, students were given the opportunity to evaluate contemporary research articles and participate in debates and discussions during their presentations. The assessment was designed to enable students to critically evaluate the knowledge they had acquired during the module, while the presentations and written assignments encouraged them to apply their newfound knowledge. **(Figure 3)**.

**Figure 2:**
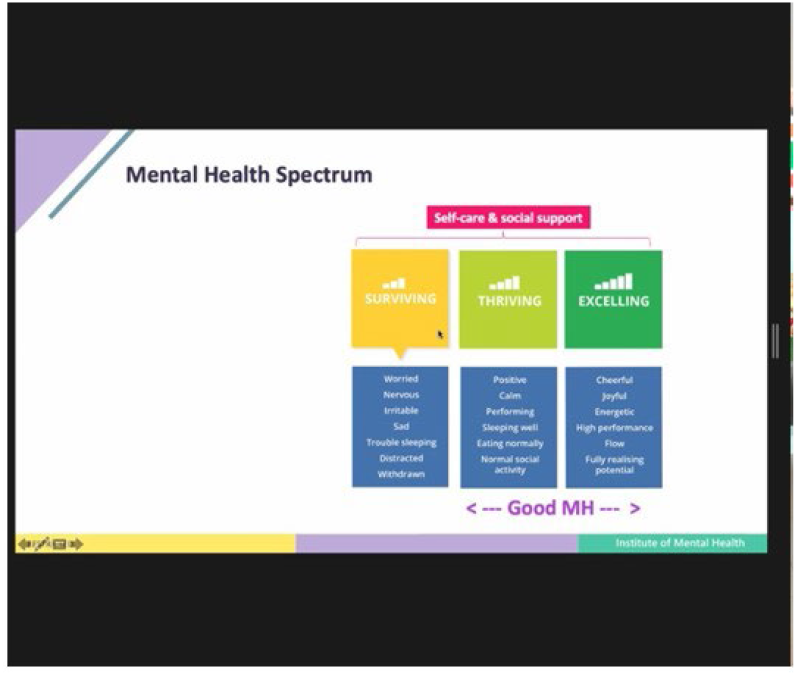
Field Visit to Institute of Mental Health, Singapore and Professional Sharing Session. A field visit to the Institute of Mental Health allows students to engage in a sharing session with professionals working in this field. Students visited the geriatric ward where early dementia patients were housed. Students learn to appreciate the operation of hospital staff caring for elderly, as well as debunk myths surrounding age-related issues. A photograph of students from Academic Year 206

**Figure 3:** Group Presentations. Students are split into groups nearer to the end of the semester, where they are tasked to critically examine an assigned research article and share with the rest of the students in the cohort. Different topics related to the issue of ageing will be discussed. A short question-and-answer session will be conducted at the end of each presentation. (Photograph)

At the end of the module, students were invited to fill in an online questionnaire to determine their opinions with regards to the 13-weeks module. This is a standard feedback exercise conducted by the University and data generated from this exercise was used in this analysis. Questions asked are broadly categorized into: (a) Overall opinion of the module; (b) Difficulty level of the module; (c) What I like about the module; and (d) What I dislike about the module. All opinions submitted were voluntary and anonymously recorded by the routine university feedback exercise. A retrospective analysis was conducted on surveys collected from all six cohorts. Qualitative reflections in this feedback exercise were considered as well.

### Survey Responses and Key Observations

Life Science Students at the National University of Singapore were offered this module on “Functional Ageing” in either their third or fourth year as part of their elective module to graduate with Bachelor’s in Science degree. Since the introduction of this module in 2016 - 2017, a total of six cohorts of students with a total strength of 268 have effectively enrolled and completed this module. Following the end of the module, students were offered to participate in a post-mortem survey questionnaire. The overall responses are collated and presented **(Figure 4-6)**. A summary of each dataset is presented in brief below.

**Figure 4:**
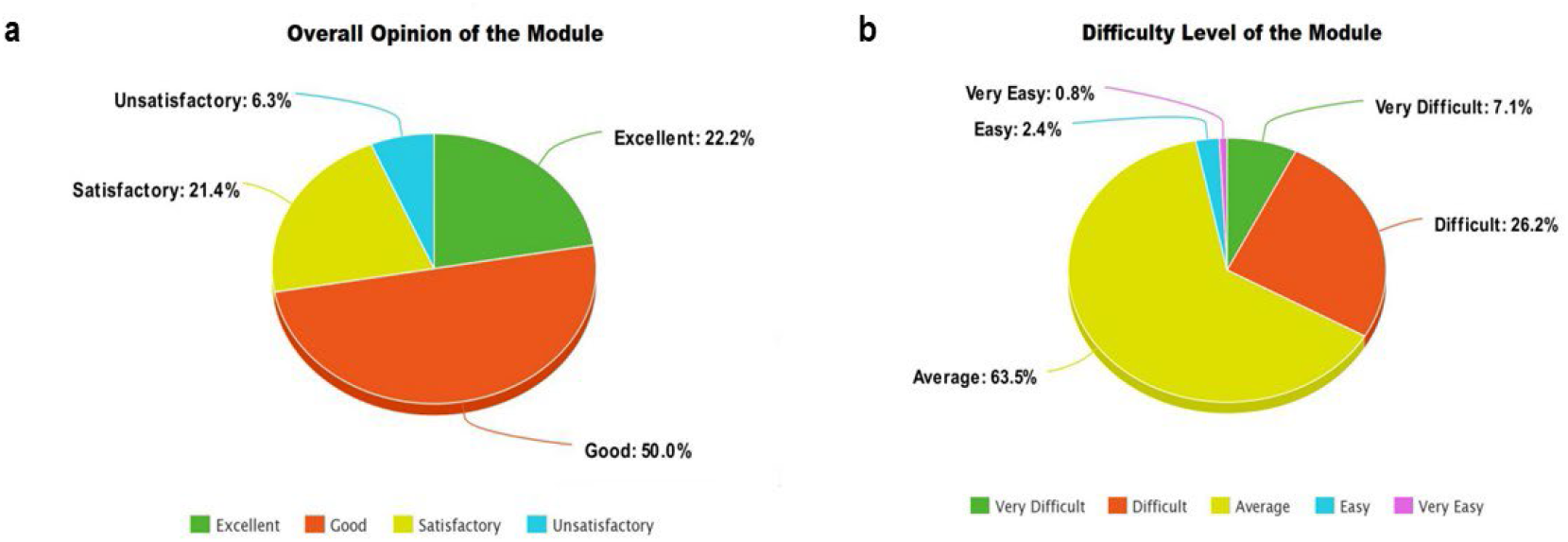
Multiple-Choice Questionnaire Responses. At the end of the module, students are invited to participate in a multiple-choice questionnaire on their (a) overall opinion as well as their (b) perceived difficulty on the module. A total of 268 students were invited for this questionnaire.

**Figure 5:**
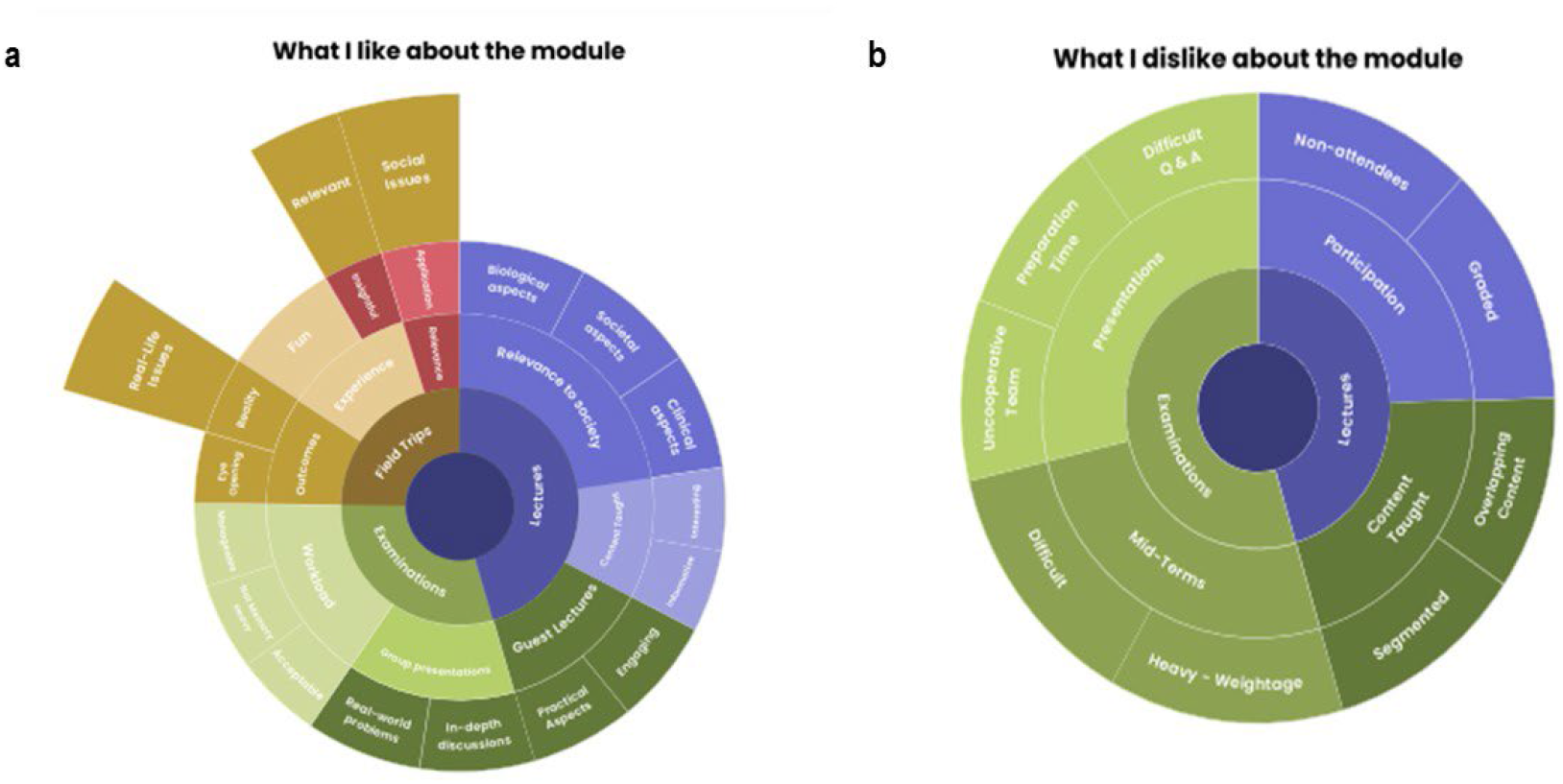
Open-Ended Questionnaire Responses. At the end of the module, students are invited to participate in an open-ended questionnaire on the reasons on (a) why they liked this module as well as (b) why they disliked this module. A total of 268 students were invited for this questionnaire.

**Figure 6:**
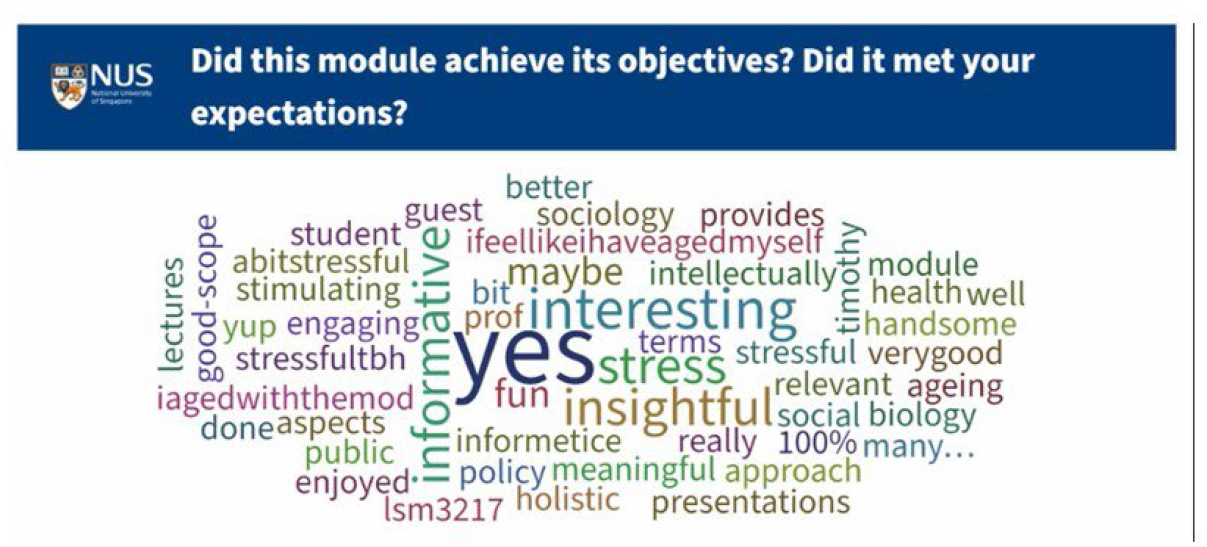
Townhall Forum. During the last lecture, students were invited to participate in a townhall forum on whether the module has met their objectives and expectations. Their anonymous responses are recorded in a mind-map.

Undergraduate students were asked to rate their overall opinion of the module, and the ratings were categorized as excellent (22.2%), good (50%), satisfactory (21.4%), and unsatisfactory (6.3%) **(Figure 4a)**. The survey results indicated that students had an overall positive experience with the module.

Students were then asked to evaluate the overall difficulty of the module, and their responses were categorized as very easy (0.8%), easy (2.4%), average (63.5%), difficult (26.2%), and very difficult (7.1%) **(Figure 4b)**. The survey results indicated that most undergraduate students found the module manageable.

Additionally, we collected open-ended responses from students when asked why they enjoyed the module. We used a sunburst chart to categorize and present the responses. The responses collected can be broadly grouped into “lectures”, “examinations”, and “field trips”. Under “lectures”, most students found the content informative and relevant to society, encompassing biological, clinical, and societal aspects. These responses align with our teaching objectives. Under “examinations”, students reported that the content tested was manageable and acceptable. Notably, many responses highlighted the usefulness of the group presentation portion of the curriculum, as it allowed for engaging in-depth discussions. Under “field trips”, students not only enjoyed themselves but also found the field trip highly relevant and valuable in developing their perspectives on ageing and societal issues, meeting our primary objective of introducing this module. **(Figure 5a)**.

Furthermore, when asked about the reasons they disliked the module, the responses were broadly categorized into “lectures” and “examinations”. Regarding lectures, some students mentioned that the content taught was fragmented and not related to each lecture, and there was feedback that the content was overlapping with other modules offered at NUS. Despite informing students that they would be assessed on their interaction/class participation, there were absentees suggesting that this mode of assessment was not a deterrent. Moving on to the examination aspect, some students found the mid-term written examinations difficult and felt that it had a heavy weightage. In addition, some students found the preparation time for the group presentation demanding and faced uncooperative team members. Finally, there were few students who found the question-and-answer session following the presentation challenging **(Figure 5b)**.

In summary, despite some challenges reported by students with regards to the academic demands of lectures and examinations, a townhall forum conducted during the final lecture revealed that this module has had a positive impact on their learning. The module covers a broad range of topics including biological, clinical, and societal aspects of ageing. Additionally, a visit to the hospital for elderly care has provided students with the opportunity to apply what they have learned in class and gain a better understanding of the societal issues caused by ageing **(Figure 6)**. This module has proven effective in equipping students with the skills to interpret ageing from different perspectives and has allowed them to appreciate the multi-faceted and complex nature of ageing. As a result, students will better appreciate the difficulties in solving age-related issues in clinical and societal contexts and strive to better understand why ageing is a persistent issue across different countries.

### Perspectives

One of the central underlying themes of the learning sciences is the inculcation of deeper knowledge in students when they engage in activities similar to everyday activities of those professionals in a discipline. At the National University of Singapore (NUS), the Functional Ageing course (from 2016 – present) implemented the STS and/or interdisciplinary teaching model, introducing cutting-edge work in aging where students learned underlying models, mechanisms, and practices applicable to various scientific disciplines. This course therefore serves to provide a “situated” view of the knowledge, where the knowledge is not just a static mental structure inside the learner’s head and instead, immerses the individual to be part of activities in which that knowledge is being applied^10^.

Module feedback exercises were conducted on five non-overlapping cohorts of students after completing the module, and their responses revealed several key findings that align with the needs of the STS training. These findings are summarised below.

#### (a) Deeper conceptual understanding

The majority of students in the cohorts expressed that they valued both memorisation and hands-on knowledge equally. While knowledge certainly matters, in the scholastic economy, memorisation of facts and various scientific pathways are not enough to fully understand the diseases in question. The module, therefore, emphasises a deeper understanding of the concepts over memorisation, as rote learning leads to mindless recitation of information. By exposing students to real-world scenarios and bringing in guest lecturers to share their practical experiences, the module provides a healthy balance in learning that creates a sense of meaning. When students develop a deeper conceptual understanding, they are able to learn facts and procedures in a much more profound and practical way that can be applied in real-world situations.

#### (b) Real-world understanding

Students who gain a deeper conceptual understanding in the classroom setting are better able to appreciate the integration of societal perspectives into the science behind ageing, not only in Singapore but also in other countries. The trips to the arts museum and the Institute of Mental Health offer students a chance to understand the ‘human touch’ that cannot be replicated in the classroom-setting. This further provides students with a wider understanding of public health issues and a more clinical understanding of various diseases.

#### (c) Building on prior knowledge

Since this module can only be taken by Year 3 and 4 students, there are overlapping concepts that build on existing knowledge. This makes students more confident in these subject areas, and students are able to articulate and reflect on this existing knowledge. This, in turn, facilitates more effective learning. Additionally, the module encourages students to reflect on and apply concepts in real-life situations, making it educationally beneficial.

Overall, The STS model of schooling is highly suitable for students pursuing careers in health care and public health research. It enables them to develop crucial knowledge and continuously enhance their understanding of the community. Nevertheless, some feedback suggested the need for greater emphasis on research techniques and molecular aspects, which relates to the importance of customized teaching methods. While it is indeed true that each student benefits differently with various teaching methods, the STS model is a crucial preliminary step towards customised learning, away from the traditional classroom-only learning pedagogy.

### Way Forward

Science education has undergone a significant change in recent decades with the incorporation of the STS approach into teaching. The traditional compartmentalized method of teaching science in theory-only has become outdated, as advancements in scientific knowledge and technology necessitate a teaching style that incorporates higher order skillsets relevant to today’s students and education system. By presenting science in an integrated fashion, students are trained to think critically, assess, and solve current issues. These skills cannot be developed if science is presented in a compartmentalised way that is disconnected from everyday reality. When presented in an integrated way, the modules become comprehensive and relevant enough for science students ^11^. The STS or interdisciplinary approach in education leads to an increase in motivation and involvement of both students and tutors, and enables educators to design more effective learning environments ^2^. Surveys report that this approach positively reinforces skills such as autonomy and critical thinking and is more motivating for students. By presenting science in a way that is relevant to everyday reality, students are better equipped to make educated decisions and develop problem-solving skills.

The STS method of learning in science has another advantage that is closely tied to the increasing importance of public health in the community. It is widely recognized that the best applicable science holds the key to solving various public health issues such as sanitation and tobacco control, as well as introducing bold policies that encourages social changes to improve health. The recent rise of COVID-19 pandemic has raised a crucial question: are science students equipped with the ability to perform rigorous research into disease risk factors, and to develop comprehensive training in advanced health disease development and management necessary to translate their scientific knowledge into public health action? As science educators, we believe that one-time science communication lectures to the public may not be impactful, instead a well-defined series of lectures in an interdisciplinary methodology especially catered to the younger generation (students) may be more effective to reach individual families.

In summary, implementing the integrated/interdisciplinary or STS method in science and medical education can lead to lifelong learning in all fields and positively impact national development. With ease of access to a variety of online resources and lectures from world renowned scholars, educators must revise their teaching pedagogy to deliver informative and not redundant content available in the public domain. Educators need to possess out-of-the box skills to facilitate efficient learning which cannot be acquired elsewhere. In our methodology presented here, students are trained to think and assess situations and apply theoretical knowledge in real-life situations, making them future ready. At a population level, an up-to-date and adaptable workforce through interdisciplinary education and training leads to increased productivity, social and institutional capital expansion, and strong impacts on economic growth.

## Acknowledgements

MPH conceived the idea, designed the syllabus and coordinated the module. MPH, GN and KM were involved in teaching the basic science component of the module. VR and SS were students of the first batch of the module (2016 – 2017) and were involved in the tutorial sessions in the subsequent years. Prof Garrie Arumugam is acknowledged for his help during the first four iterations of this module. We thank all the guest lecturers of this module, Drs Lydia Seong, Resham Lal Gurung, Feng Lei,, Rahul Malhotra, Chetna Malhotra, Stella Schwe. Special thanks to Professor Leonard Hayflick, Department of Anatomy, University of California San Francisco, USA and Professor Kua Ee Hock, Department of Psychological Medicine, National University of Singapore for their special lectures. Institute of Mental Health and other elder care centers for hosting the students for their field visits. Mr Karthik Prathaban and Dr Varsha Hande are thanked for critical reading of the manuscript.

## References

1 Clancey, G. K. & Chhem, R. K. Radiation risk communication in Fukushima from a STS perspective. (2023).

2 Primastuti, M. & Atun, S. Science Technology Society (STS) learning approach: an effort to improve students’ learning outcomes. Journal of Physics: Conference Series 1097, 012062, doi:10.1088/1742-6596/1097/1/012062 (2018).

3 Benjamin, K. A. & McLean, S. Change the medium, change the message: creativity is key to battle misinformation. Adv Physiol Educ 46, 259–267, doi:10.1152/advan.00021.2021 (2022).

4 Houston, J. B., Hansen, G. J. & Nisbett, G. S. Influence of User Comments on Perceptions of Media Bias and Third-Person Effect in Online News. Electronic News 5, 79–92, doi:10.1177/1931243111407618 (2011).

5 Sicherl-Kafol, B., & Denac, O.. The importance of interdisciplinary planning of the learning process. Procedia - Social and Behavioral Sciences 2 (2). doi:10.1016/j.sbspro.2010.03.752 (2010).

6 Sargent-Cox, K. Ageism: we are our own worst enemy. Int Psychogeriatr 29, 1–8, doi:10.1017/S1041610216001939 (2017).

7 WorldHealthOrganisation. Ageing and health, <https://www.who.int/news-room/fact-sheets/detail/ageing-and-health> (2022).

8 Goh, O. Successful Ageing — A Review of Singapore’s Policy Approaches. (Civil Service College, Singapore, https://www.csc.gov.sg/articles/successful-ageing-a-review-of-singapore%27s-policy-approaches, 2006).

9 Hande, V., Prathaban, K. & Hande, M. P. Educational dialogue on public perception of nuclear radiation. Int J Radiat Biol 98, 158–172, doi:10.1080/09553002.2022.2009147 (2022).

10 Greeno, J. G. in The Cambridge handbook of: The learning sciences. 79–96 (Cambridge University Press, 2006).

11 Ferreira, S. & Morais, A. M. The Nature of Science in Science Curricula: Methods and concepts of analysis. International Journal of Science Education 35, 2670–2691, doi:10.1080/09500693.2011.621982 (2013).

